# Universal Spectrum Explorer: A standalone (web-)application for cross-resource spectrum comparison

**DOI:** 10.1101/2020.09.08.287557

**Authors:** Tobias Schmidt, Patroklos Samaras, Viktoria Dorfer, Christian Panse, Tobias Kockmann, Leon Bichmann, Bart van Puyvelde, Yasset Perez-Riverol, Eric W. Deutsch, Bernhard Kuster, Mathias Wilhelm

## Abstract

Here we present the Universal Spectrum Explorer (USE), a web-based tool based on IPSA for cross-resource (peptide) spectrum visualization and comparison (https://www.proteomicsdb.org/use/). Mass spectra under investigation can either be provided manually by the user (table format), or automatically retrieved from online repositories supporting access to spectral data via the universal spectrum identifier (USI), or requested from other resources and services implementing a newly designed REST interface. As a proof of principle, we implemented such an interface in ProteomicsDB thereby allowing the retrieval of spectra acquired within the ProteomeTools project or real-time prediction of tandem mass spectra from the deep learning framework Prosit. Annotated mirror spectrum plots can be exported from the USE as editable scalable high quality vector graphics. The USE was designed and implemented with minimal external dependencies allowing local usage and integration into other websites (https://github.com/kusterlab/universal_spectrum_explorer).

## Introduction

In mass spectrometry (MS), peptide spectrum matches (PSMs) of tandem mass spectra (MSMS) play a fundamental role independent of the acquisition method or analysis. PSMs most commonly originate from data dependent acquisition (DDA) experiments, where MSMS spectra are matched against in silico generated fragmentation patterns(1)For targeted (2) and data independent acquisition (3, 4) proteomics, previously generated PSMs are (often) a prerequisite for the confident detection and quantification of peptides. In contrast to DDA experiments, the incorporation of fragment intensities plays a crucial role here in order to increase the specificity of the detection.

Being the foundation for (almost) all proteomic experiments, the correctness of PSMs is of utmost importance. This is particularly the case when e.g. single amino acid permutations(5), alternative reading frames(6) or novel HLA peptides(7) are identified. Although methods exist to control the global and local false discovery rate(8, 9), the absolute number of false positive identifications will increase according to the size of the dataset (10, 11). Furthermore, previous examples have shown that false positives can accumulate and surpass (global) FDR criteria for certain classes of peptide when not controlled for specifically(12).

Over the last years, the number of available (reference) spectra in public repositories(13–15), synthetic resources(16) and high quality predictions(17–20) has increased drastically. In principle, this opens up the possibility to manually verify PSMs by comparing an experimental spectrum to a reference spectrum. However, two challenges remain. First, the retrieval of reference from the three mentioned sources is often cumbersome or requires expert knowledge. The HUPO-PSI addressed parts of this recently by specifying the Universal Spectrum Identifier (USI, http://www.psidev.info/usi), describing a virtual path to locate a spectrum and a possible interpretation in public repositories. However, other sources of spectra such as prediction services cannot be referenced by this identifier. Second, the generation and scoring of mirror spectra currently requires the installation of complete software suites such as Skyline(21, 22), Trans-Proteomic Pipeline(23) or OpenMS(24).

Here, we present the Universal Spectrum Explorer (USE, https://www.proteomicsdb.org/use), a versatile web-based tool, that allows the visualization and comparison of any two spectra. The USE is based on the Interactive Peptide Spectrum Annotator (25) and was implemented in part at the EuBIC-MS Developer’s Meeting 2020 (26). In the USE, each spectrum can be entered in three different ways. First, originating from a local source (i.e. spectrum extracted from a raw file) by uploading a peak list. Second, by providing the USI of a spectrum available in a public repository allowing users to access more than 470 million spectra in PRIDE alone. Third, by retrieval of spectral information from a newly defined and developed REST(27) interface for resources that are currently not covered by the USI schemata. An example implementation of this is provided on ProteomicsDB(28, 29) and allows access to either spectra acquired in the ProteomeTools(6) project (more than 5 million spectra) or real-time predicted spectra by accessing our deep-learning framework Prosit(17). The settings of the USE are open to the users to interact with the spectra, allowing adjusting and zooming into spectra as well as highlighting of matched peaks. If the user is satisfied with the visualization, we offer a download in tabular format for reprocessing or an export of the figures as editable and scalable high quality vector graphics. The USE was developed with minimal external dependencies and can be integrated into third party website. The source code of is available on Github (github.com/kusterlab/universal_spectrum_explorer).

## Experimental Section

### Software implementation

The code base of USE is separated into three independent packages. The proteomicsJS package provides first basic algorithms and functionalities to work with spectral data in Javascript. It has no special requirements, can be used in any Javascript framework and is accessible on Github (https://github.com/kusterlab/proteomicsJS).

The USE (https://github.com/kusterlab/universal_spectrum_explorer) is the client-facing interactive web visualizer using AngularJS based on IPSA. Its sole purpose is to provide a user interface and leverages proteomicsJS for all use cases.

The USE_embedded (https://github.com/kusterlab/use_embedded) is a static executable available for all major operating systems (Mac, Windows and Linux) and architectures (ARM, x86 and PowerPC). It has two major use cases. First, it combines proteomicsJS as well as the USE to a single file and allows the USE to be easily distributed either on local machines or any server. Second, it can be used as a transparent proxy that handles requests to all Proteomics eXpression Interfaces (PROXI) (30) of all repositories. This might be necessary when the USE is hosted on a server using Transport Layer Security (TLS) encryption, .e.g. https://proteomicsdb.org/use, because current security policies do not allow to query non-TLS endpoints from existing TLS connections.

### Peak matching

The peak matching algorithm applied by USE is designed towards high-resolution, accurate-mass (HRAM) data by exactly matching peaks between two spectra according to their mass to charge ratio (m/z). Matched peaks must adhere to relative or absolute matching tolerance defined in ppm or Dalton. Before spectra are submitted to the matching procedure, the intensities of isobaric fragments are summed up. This may happened particularly when predicted spectra are used for comparison (e.g. Prosit). For example, the predicted fragmentation pattern of the doubly charged peptide AEAEAQAEAAR is predicted to have 0.77% and 62.5% relative intensity for the isobaric fragments b2+ (sequence AE) and b4(2+) (sequence AEAE), respectively, at a collision energy of 30 by Prosit. The matching procedure is outlined below:
INPUT: two arrays, named S_1_ and S_2_, of (m/z, intensity, peak_id) triplets sorted by m/z OUTPUT: one array S_matched_ of elements of m/z_1_, intensity_1_, peak_id_1_, m/z_2_, intensity_2_, peak_id_2_. Peaks of S_1_ that couldn’t be matched to a peak of S_2_ are included in the array as well, but the m/z_2_, intensity_2_, peak_id_2_ is set to NA. The same is applied to peaks of S_2_. A peak can just occur once.

1. For every peak in S_1_, search for the closest peak in the S_2_ according to matching tolerances

a. If multiple peaks in a queried spectrum match to a reference spectrum, select the most intense one and unassign the rest
2. Repeat step 1 but search for every peak in S_2_ a matching partner in S_1_
3. Concatenate the results of step 1 and 2 and remove duplicate peak pairings. In case of contradictory assignments of peaks, select the one with the most summed intensity.

### REST interface specification

The USE can receive spectral data from any web endpoint adhering to a minimal schema. The USE sends a request as a JSON encoded list of sequence, collision energy (CE), charge and modification strings. If a peptide is modified at multiple residues, the modifications are appended to each other and separated by comma. The expected output is mirroring the input by also being a JSON encoded list of sequence, precursor charge, collision energy as well as a list of ions (m/z, intensity). A complete description can be examined at https://github.com/kusterlab/universal_spectrum_explorer/wiki.

## Results and Discussion

### Design of USE

The newly developed Universal Spectrum Explorer (USE) is based on the web-based spectral analysis tool IPSA(25). The main scope of the USE is the comparison of any two spectra while maintaining most of the functionality originally implemented. For this, we optimized all core features (spectrum comparison and annotation) to be easily handled by modern browsers and, as long as no spectra from external resources are requested, the USE can be used entirely offline (locally). To ensure this, some functionalities of IPSA were re-implemented to support local execution. In addition, all functionalities were developed without the use of additional external libraries, minimizing dependencies and overhead. This not only enables future contributors to easily and quickly understand and extend the current functionalities but also minimizes required changes in the codebase resulting from changes in external libraries. As a result, no special software is required to execute the USE and can be opened directly with any browser. During development, we followed the test-driven development principles and achieved close to 80% code coverage.

For the comparison of any two spectra, the user is asked to define the source or spectral data of the top and bottom spectrum by either providing the spectral information directly, a USI, or information necessary to retrieve a spectrum from any other source implementing the newly developed interface. By this, the USE offers the functionality to allow the comparison of, for example, a locally acquired spectrum to a spectrum stored in PRIDE, a modified to an unmodified peptide spectrum, two spectra of the same peptide acquired at different collision energies or two in silico generated spectra from two different sources.

### Data import and annotation

The simplest way to import data into the USE is the manual input tab where a user specifies general metadata such as the unmodified sequence and the precursor charge. The webform for the spectral data can be filled directly by pasting data from any spreadsheet program (Google sheets/excel), a csv-file or the output of Xcalibur Software of Thermo Fisher Scientific. Post-translational or other peptide modification can be either selected from a predefined dropdown menu or added manually by the user. Manually added modifications are stored locally and will neither be persistent nor available to other users (Figure 1a).

**Figure 1.**
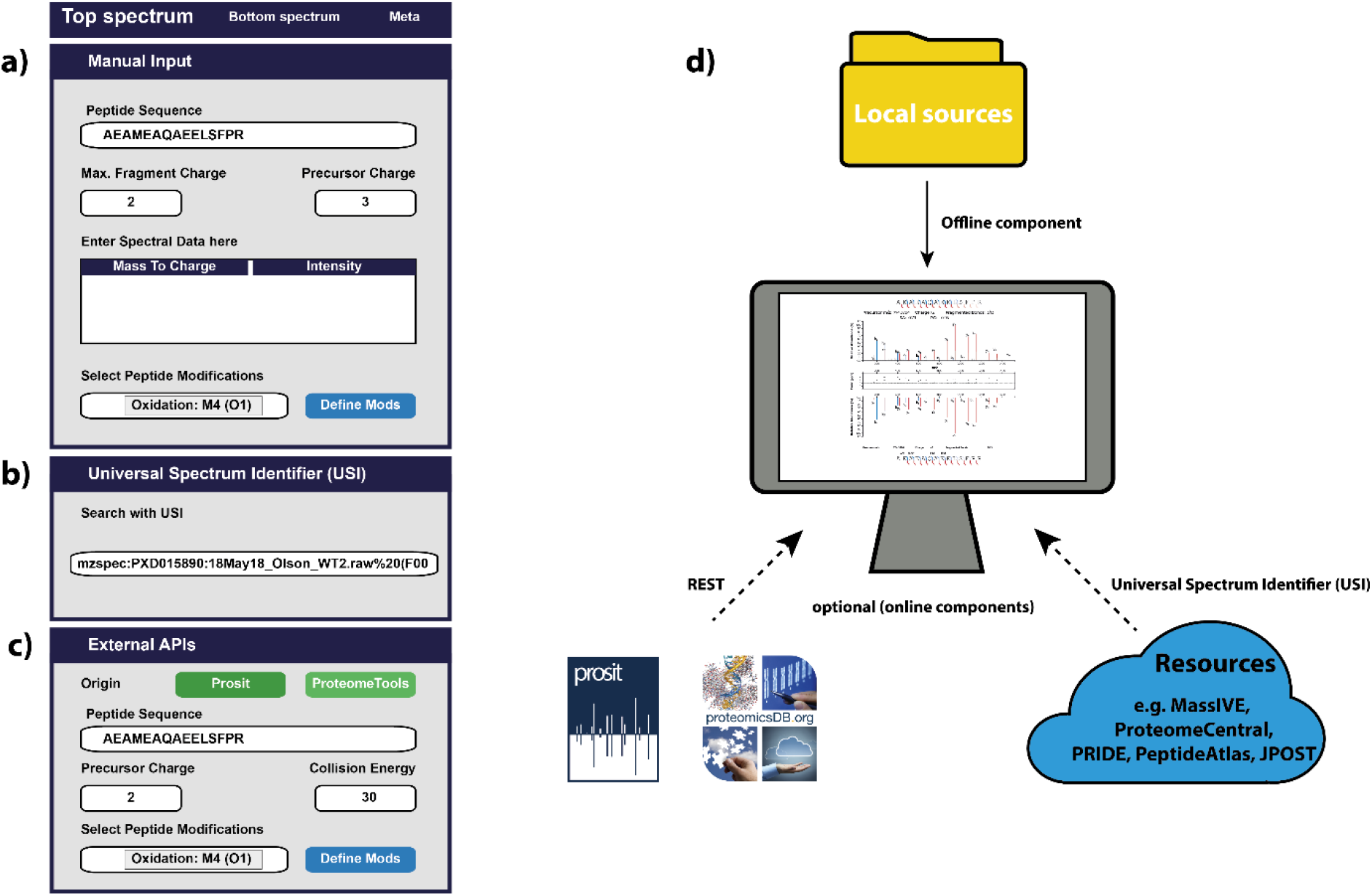
The USE interface. a) The “manual input” section allows the definition of every setting by hand up to the manual generation of modifications. b) The “USI” section allows pasting of an USI and automated retrieval from three different data repositories implementing the USI standard. c) The “external APIs” section is an extension of the USI and allows a retrieval of spectral data from any resource implementing our defined REST endpoint. ProteomicsDB offers additional endpoints and provides data from the ProteomeTools project and the peptide fragmentation prediction tool Prosit. d) The input of spectral data for the USE can originate from various sources, such as local mass spectrometer files, public data repositories and prediction services.

A second option is the retrieval of spectra from public resources by supplying a USI (Figure 1b). The USE will then attempt to retrieve the spectrum using the Proteomics eXpression Interfaces (PROXI)(30) interface from many resources within the ProteomeXchange(31)(PeptideAtlas(32), MassIVE (33), PRIDE(15), JPOST(34)). The peptide sequence and present modifications are parsed based on the ProForma specification (35) within the USI. The peptide sequence, precursor charge and list of extracted modifications are directly submitted to the USE. The spectrum is then inserted into the manual input which ensures a consistent processing of the data. If a modification cannot be parsed correctly from the USI, the user is alerted and able to add the modification manually.

Lastly, a new interface was defined allowing access to other resources not adhering to the USI standard (yet). This interface was implemented in ProteomicsDB allowing access to spectra acquired in the ProteomeTools project and direct access to Prosit for real-time spectrum prediction (limited currently to the monoisotopic y/b ions generated by HCD fragmentation). Similar to the manual user input tab, the peptide sequence, precursor charge and modifications are specified. In addition, the collision energy must be provided. Since the ProteomeTools resource might not (necessarily) contain the supplied CE, the spectrum with the closest matching collision energy is automatically selected (Figure 1c). If the data is requested from Prosit or ProteomeTools, similarly to the USI tab (Figure 1d), the metadata and spectral information is stored in the manual input for consistent spectral processing.

The original IPSA viewer focused on the comprehensive annotation of fragment peaks in a spectrum by utilizing a remote server for computation. We re-implemented the core annotation functions of the IPSA backend in JavaScript in order to minimize external dependencies. The annotation allows the highlighting of peaks in the spectrum viewer given a selection of fragment types and annotation thresholds. All necessary parameters for annotation and visualization can be set in the metadata section of the viewer.

### Spectrum similarity calculation

In order to assess the similarity of two spectra, peaks occurring in either spectrum need to be matched to the other. The process of matching peaks for two spectra does not require the annotation of the peaks and the result of this process is referred to as a “matched spectra” in this manuscript (see Experimental Section).

With matched spectra at hand, the similarity of two spectra can be scored in two ways: First, the similarity can refer to the question “How identical are these two spectra?”. This metric will take all peaks (matched or unmatched) of both spectra into account. Peaks missing in one or the other spectrum will penalize the similarity score (Figure 2). However, this can lead to (somewhat artificially) low similarity scores, when for example two spectra are compared where one contains a significant co-isolation. In such cases, a different spectral similarity score might be useful. This addresses the question: “How well is one spectrum contained in another?”. For this purpose, all peaks exclusively present in one spectrum are discarded prior to the similarity calculation (Figure 2). This ensures that unmatched peaks from one spectrum do not penalize the similarity artificially, e.g. excluding peaks from co-isolated peptides. Because the top spectrum can be contained in the bottom spectrum, and vice versa, the USE displays all three possible spectral similarities. Pearson’s correlation coefficient (PCC) and (normalized) spectral (contrast) angle (SA) are used as similarity measures in the USE. Although both measures correlate well, we(17) and others(36) have previously observed that the spectral angle offers higher sensitivity to differentiate similar spectra, while most users will be more familiar with PCC.

**Figure 2.**
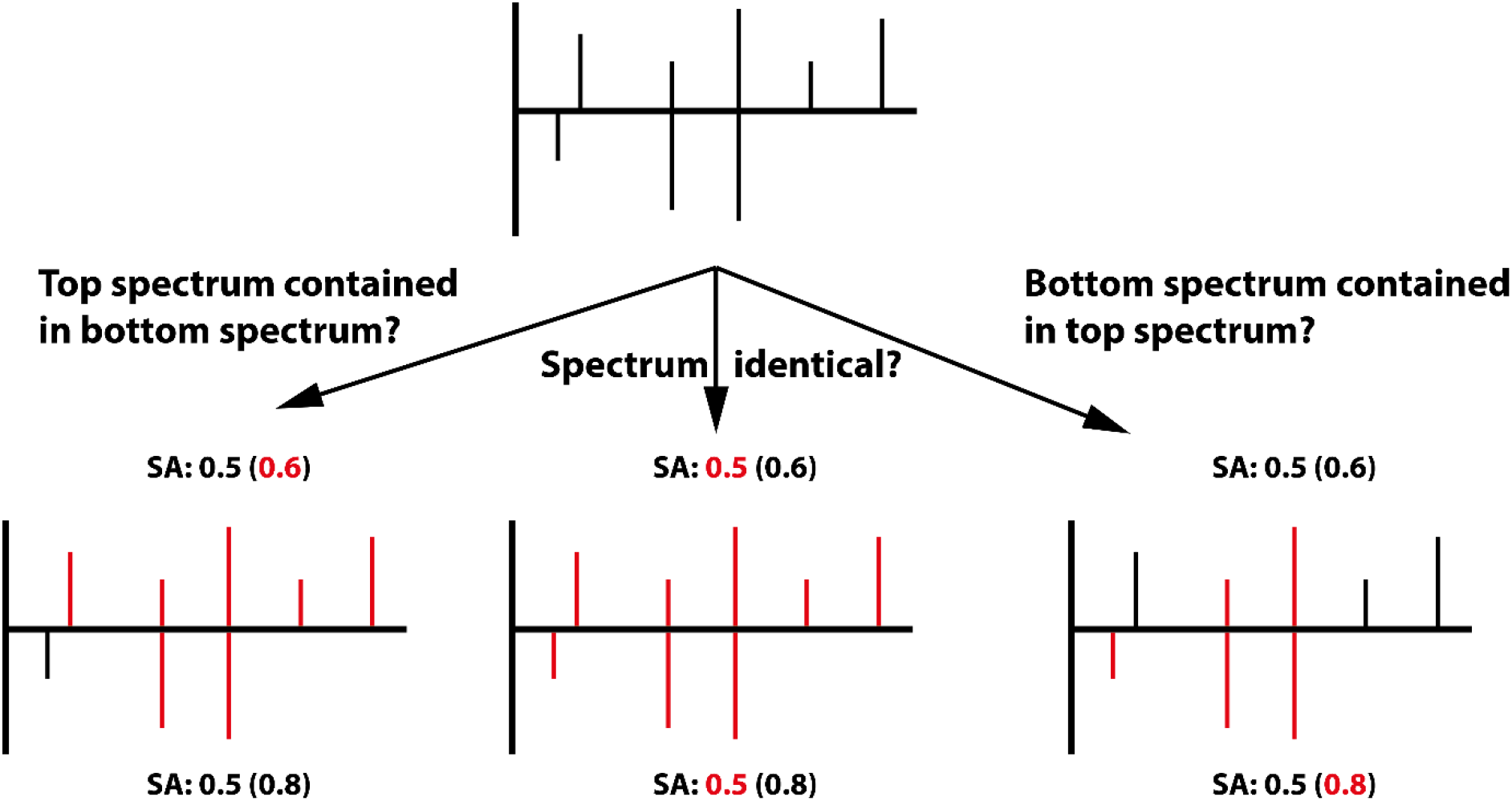
Spectral similarity between two spectra (top mirror plot) can be used to address three questions: 1) Whether two spectra are identical (bottom middle), using all peaks for similarity calculation, 2) whether the top spectrum is contained in the bottom spectrum (bottom left), discarding unmatched peaks in the bottom spectrum, or 3) whether the bottom spectrum is contained in the top spectrum (bottom right), discarding unmatched peaks in the top spectrum. The spectral angle (SA) is highlighted in red for each of the three cases. Peaks used to calculate the spectra angle are highlighted in red for each case.

The USE generates an interactive view of the spectrum comparison (Figure 3), consisting of two spectrum visualizations (Figure 3a and c) showing both selected spectral data and a peak-assignment/intensity difference plot in the middle (Figure 3b). All three plots are coupled to each other, allowing zooming on the m/z axis as well as hovering over peaks or error circles highlighting the matched peak in the other spectrum. The spectrum plots show the peptide sequence, detected fragments and precursor information as originally implemented in the IPSA viewer. In addition, we show the similarity measures PCC and SA resulting from the three different strategies when merging spectra (see Experimental Procedures). The peak-assignment/intensity difference plot in between both spectra visualizes the difference of the selected spectra by showing matched peaks in black and unmatched peaks in grey dots. The size of the dots correlates with the difference in intensity whereas the height of the dots represents the observed mass difference in part per million (0 for unmatched peaks). Note, the mass difference is calculated with respect to the average *m*/*z* of the two matched peaks. This is to avoid having none or multiple mass differences when either the theoretical mass is unknown (i.e. comparison of two experimental spectra where matching peaks can have no annotation) or the theoretical masses of the matched peaks differ (i.e. matching peaks but different annotation due to the selection of two different peptide sequences for top and bottom), respectively. While this will not represent the real mass errors, the overall trend will still be conserved. More importantly, this allows a generic design and implementation of the USE where no assumptions are made as to which spectrum is the experimental or reference spectrum and the intensity difference plot will represent a matching peak pair with one dot.

**Figure 3.**
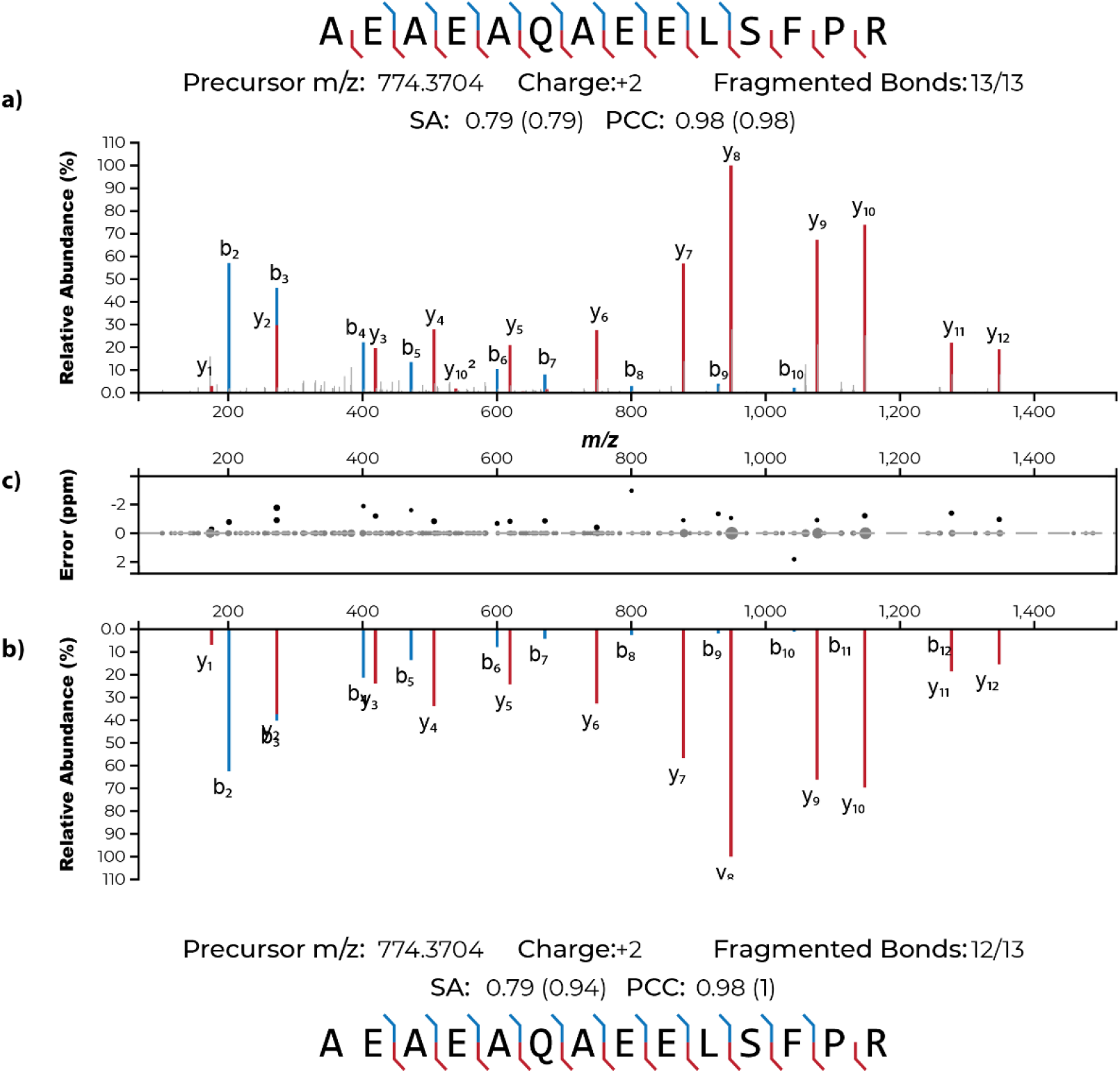
Mirror spectrum visualization of the USE. The top spectrum representing the doubly charged peptide spectrum match AEAEAQAEELSFPR (PXD015890) (USI: mzspec:PXD015890:18May18_Olson_ WT2.raw%20(F001551).mzid_ 18May18_Olson_ WT2.raw_(F001551).MG F:index:6913:AEAEAQAEELSFPR/2 stored at PRIDE), the bottom spectrum a predicted spectra by Prosit of the same sequence and precursor charge. a) +b) MSMS spectrum visualization. Annotated peaks are coloured according to their inferred ion type (blue = b series, red = y series; colouring can be adjusted via the settings tab). Underneath the bottom spectrum as well as above the top spectrum meta information is shown. Spectral angle (SA) and Pearson’s correlation coefficient (PCC) describe the similarity of the two spectra. Similarity measures in brackets over the top spectrum describe how similar both spectra are based solely on the peaks of the top spectrum. The same is shown for the bottom spectrum based solely on the peaks of the bottom spectrum. c) Visualization of the mass and intensity discrepancy between peaks in both spectra. Black dots represent matched peaks and their mass error. Grey dots depict unmatched peaks. The size of the dots is used to indicate the intensity discrepancy of individual peaks.

## Conclusion

Here we presented the Universal Spectrum Explorer (USE), an open-source and web-based tool for the comparison of any two spectra. It allows the quick and easy cross-resource comparison of two spectra. We strongly believe that the USE will become an important tool for researchers and reviewers to be able to directly and quickly investigate, compare and validate spectra and PSMs of claimed peptide identifications. This circumvents the necessity to install additional third party software and to download large amounts of raw data. With the ability to directly link to spectra stored in public repositories by using the USI, we envision that shareable links can be provided as part of the Supplementary Information of a manuscript (e.g. https://www.proteomicsdb.org/use/?usi=mzspec:PXD000561:Adult_Frontalcortex_bRP_Elite_85_f09:scan:17555:VLHPLEGAVVIIFK/2). For the visualization of single spectra or unidentified spectra, we recommend to use IPSA(25) or the Metabolomics Spectrum Identifier Resolver (37).

Furthermore, the modular layout of the USE allows complete or partial integration into third party websites. Due to its minimal dependencies, the USE requires no complex built system and can also be run locally. In order to allow researchers to benefit from recent advances in spectrum predictions using e.g. deep learning, we developed a new REST interface allowing the retrieval of spectral data from other sources. We showcase this in ProteomicsDB which enables the USE to retrieve predicted spectra from Prosit in real-time. Any other prediction service can in principle be integrated by this mechanism and would allow scientists to pick and evaluate which prediction engine fits their use case best.

Assessing the global probability or likelihood of a correct match by comparing an experimental (or query) spectrum to a reference spectrum still remains difficult. Although spectral similarity measures such as PCC and SA specifically take fragment intensity into account, the definition of a threshold indicative of spectral identify requires the estimation of a similarity distribution(38) or scoring of all possible spectral combinations (39). Both methods are, compared to the classical target-decoy estimation, computationally expensive and many alternative amino acid combinations can be found whose spectra only vary to a very minor degree. The USE and particularly the implemented interfaces to public repositories and prediction servers might play an important role in the development of robust methods estimating such thresholds.

We firmly believe that the USE will become a useful tool in many areas of research. For example, the comparison of spectra of modified and unmodified counterpart peptides helped to identify modification specific diagnostic ions(40). The USE could be used initially to investigate such differences. Similarly, the USE can be used to investigate (and if necessary, even manually adjust) peptide identification results retrieved from de novo search engines. Since reference spectra of peptide sequences suggested by de novo search engines are likely not present in public repositories, the integration of fragment intensity predictions in the USE offers a unique way to judge and potentially validate such PSMs. Due to Prosit’s unique ability to predict spectra at various collision energies, the USE could be used to manually optimize the collision energy used for targeted experiments in order to maximize the intensity of particular fragment ions.

## Abbreviations

API: Application Programming Interface
CE: Collision Energy
DDA: Data-dependent Acquisition
DIA: Data-independent acquisition
EuBIC: European Bioinformatics Community
FDR: False Discovery Rate
HCD: Higher-energy Collision Dissociation
HLA: Human Leukocyte Antigen
HRAM: High-Resolution and Accurate Mass
IPSA: Interactive Peptide Spectral Annotator
MS: Mass Spectrum
MSMS: Tandem Mass Spectra
NPM: Node Package Manager
PCC: Pearson’s Correlation Coefficient
PRM: Parallel Reaction Monitoring
ProForma: Proteoform and Peptidoform Notation
PROXI: Proteomics eXpression Interfaces
PSM: Peptide Spectrum Match
PTM: Post-Translational Modification
REST: Representational State Transfer
SA: Spectral Angle
TLS: Transport Layer Security
USE: Universal Spectrum Explorer
USI: Universal Spectrum Identifier

## Acknowledgements

We are thankful for EuBIC-MS for organizing the EuBIC-MS developer meeting 2020. Our tool is heavily influenced by the keynote talks and the initial prototype was developed as part of the hackathon. The authors wish to also thank all members of the Kuster laboratory. This work was in part funded by the German Federal Ministry of Education and Research (BMBF; grant No 031L0008A and 031L0168) and the European Union’s Horizon 2020 Program under Grant Agreement 823839 (H2020-INFRAIA-2018-1; EPIC-XS). The IBM infrastructure hosting ProteomicsDB and Prosit is operated and maintained by the SAP University Competence Center (UCC) of the Technical University Munich.

## Code availability

All source code has been made available on GitHub

- https://github.com/kusterlab/universal_spectrum_explorer
- https://github.com/kusterlab/use_embedded
- https://github.com/kusterlab/proteomicsJS

## For TOC only

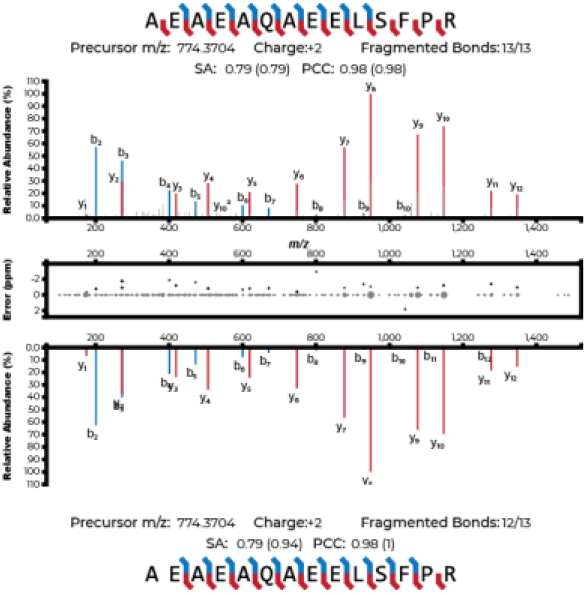

